# Targeted fusion of Antibody-Secreting Cells: unlocking monoclonal antibody production with hybridoma technology

**DOI:** 10.1101/2025.03.19.644155

**Authors:** Fanny Rousseau, Catherine Menier, Patricia Brochard, Stéphanie Simon, Karla Perez-Toralla, Anne Wijkhuisen

**Affiliations:** Université Paris-Saclay, CEA, INRAE, Médicaments et Technologies (MTS), 91191 Gif-sur-Yvette, France

**Author notes:** These authors contributed equally to this work (Karla Perez-Toralla, Anne Wijkhuisen). **<u>Corresponding author:</u>** Anne Wijkhuisen. Email address, Tel: +331, 69 08 80 35.

**Keywords:** Hybridoma technology, Monoclonal antibodies, Antibody-secreting cells, Cell fusion, FACS

## Abstract

Monoclonal antibodies (mAbs) produced by hybridoma technology have extensively proved their value for therapeutic, diagnostic, and biomedical research applications, despite the reported low fusion yields between short-lived B cells and immortal myeloma cells. To improve the efficiency of this process and accelerate the development of new mAbs, we characterized and isolated antibody-secreting cells (ASCs) from the spleen of immunized mice before cell fusion. This approach resulted in a high yield of hybridoma generation by increasing the probability of successive pairing between the most suitable cell fusion partners. Specifically, we developed an optimized workflow combining Fluorescence-Activated Cell Sorting (FACS) with antibody secretion assays, using a panel of five cell-surface markers (CD3, TACI, CD138, MHC-II, and B220) that allowed us to identify a particular ASC subset with key characteristics. Such ASCs exhibited a plasmablast phenotype with high MHC-II expression and secreted high levels of Ag-specific antibodies in immunized mice. These features were also found in hybridomas, suggesting a preferential fusion of myeloma cells with this ASCs subset. Finally, the targeted electrofusion of TACI^high^CD138^high^ sorted ASCs led to a 100% fusion yield compared to a non-targeted approach. In particular, over 60% of these generated hybridomas secreted Ag-specific mAbs. Collectively, these results pave the way for a highly efficient method to produce new mAbs by cell fusion, which could facilitate hybridoma generation and expand therapeutic applications of mAbs.

## Introduction

Antibodies are large glycoproteins also known as immunoglobulins (Ig) which are produced, cell-surface expressed and secreted by B lymphocytes in response to an infection or immunization. Each mature B cell only produces one type of antibody with the same single antigenic specificity. These monoclonal antibodies (mAbs), highly specific and affine, have established themselves as one of the largest groups of biologics (biotherapeutic proteins) now used in various therapeutic applications (1,2). Since the approval of the first therapeutic mAb, the muromonab-CD3, in 1986, the Food and Drug Administration (FDA) has approved more than 100 therapeutic mAbs for various diseases, including autoimmune disorders and inflammatory diseases, and nearly half for cancer (3). In addition to their therapeutic potential, mAbs are widely used as *in vitro* and *in vivo* diagnostic tools due to their high specificity and targeted binding affinity. In medical imaging, mAbs or their derivatives are also tools to detect tumors in early stages of development (4). In research, antibodies are essential in many techniques, such as ELISA, flow cytometry, immunocytochemistry and immunohistochemistry, immunofluorescence, Western blotting, immunoprecipitation, and more (5,6). The increasing importance of mAbs in therapeutics and diagnostics highlights the critical need to accelerate and optimize their production and development methods.

As early as 1975, Köhler and Milstein proposed hybridoma technology to immortalize B cells producing mAbs since these cells cannot survive in culture for more than a few days (7). Hybridomas are cells formed by fusion between a short-lived antibody-producing B cell and an immortal myeloma cell. This fusion uses spleen cells harvested after mouse immunization with the antigen (Ag) of interest, generally using polyethylene glycol (PEG). After a few days of culture in a selective culture medium that allows only fused cells to survive, Ag-specific antibodies are evaluated in hybridoma culture supernatants. Cells secreting Ag-specific antibodies are subsequently cloned, usually by limiting dilution, to isolate single hybridomas secreting Ag-specific mAbs and achieve monoclonality. Each hybridoma clone constitutively produces a large amount of one specific mAb, and cell lines are cryopreserved to ensure long-lasting mAb production. Currently, more than 90% of FDA-approved therapeutic mAbs are generated by hybridoma technology (8). These mAbs being generated mostly from mice, chimerization, and humanization techniques were rapidly developed to limit their immunogenicity and improve their efficacy.

The main advantage of hybridoma technology relies on using B cells from mice, preserving the natural ability of the murine immune system to generate highly specific mAbs. Indeed, the B cells that enter the cell fusion process to produce Ab-secreting hybridomas have undergone all steps of the differentiation process, such as somatic hypermutations in the complementary-determining regions (CDRs) of the variable genes, thus leading to a strong increase in affinity for the antigen. This differentiation process occurs in the secondary lymphoid organs, in the germinal centers formed during an immune response, as a result of follicular helper T and activated B cells collaboration, leading to plasmablasts and plasma cells that secrete a large amount of Ag-specific antibodies (9–11).

Despite its robustness, one major limitation of the hybridoma technology is the rarity of the fusion events between B cells and myeloma cells (5 x10^−6^ efficiency with conventional PEG-based fusion (12)), resulting from the random paring between the two cell partners and the low efficiency of the PEG-based cell fusion mechanism. Consequently, many B cells that may produce high-affinity antibodies to the target antigen are lost. To address this challenge, strategies such as B cell enrichment prior to cell fusion have been employed to facilitate successive pairing, thereby enhancing cell fusion efficiency (13,14). In addition to PEG-based fusion methods, alternative approaches, including electrofusion, have also been investigated to enable the generation of stable antibody-producing hybridomas. This technique uses electrical pulses to induce membrane fusion, allowing for greater control and improved hybridoma yield (15).

Other methods for producing antibodies directly from B cells have evolved as alternatives to cell fusion. As early as 1995, Lagerkvist et al. isolated single, Ag-specific human B cells using Ag-coated magnetic beads to generate Ag-specific recombinant antibody fragments (16). O Starkie et al. developed a multi-parameter flow cytometry single, IgG^+^ memory B cells sorting technique, followed by direct amplification of V_H_ and V_L_ region encoding genes and subsequent expression in cell culture systems, allowing the preservation of the original V_H_ and V_L_ pairing (17). It is undeniable that single-cell recombinant antibody technologies offer the essential advantage of long-term stability of transfected cell lines compared to hybridomas. Following the rearrangement of their genetic material, hybridomas can, in some cases, contain one or more additional productive heavy or light chains, which can induce the risk of degrading some properties of the antibodies such as their specificity (18,19). However, single B cell-based techniques remain time-consuming and challenging, requiring many micromanipulations and further development.

Despite its limitations, hybridoma technology remains robust and requires no specific instrumentation, making it easy to implement in research laboratories. In particular, we have developed numerous mAbs for immunological tests for the diagnosis/detection of infectious diseases or biological warfare agents (including commercially available tests) and immunotherapies (20,21).

To improve the hybridoma technology further, we propose a new method based on the selection of antibody-secreting cells (ASCs), the terminally differentiated form of B cell, before the cell fusion step. Specifically, by selecting ASCs, including plasmablasts and plasma cells, based on cell-surface markers expression, the probability of successive fusion between both cell fusion partners of interest can be increased, leading to a high proportion of functional hybridomas among all fused cells (high yield).

First, we implemented an optimized labeling panel and workflow to characterize splenic cells using flow cytometry and cell sorting. To identify the cell populations of interest before fusion, we measured their Ab secretion levels in culture supernatant several days after sorting, using dedicated ELISA assays. We compared the results from naive and immunized mice to highlight the differentiation process of B cells and identify the activated cell populations following immunization. We then applied the developed labeling panel to analyze a hybridoma cell mixture obtained by conventional fusion (before culture and cloning). We analyzed the expression of ASC cell-surface markers and the potential of the different sorted populations to secrete Ag-specific antibodies. Finally, we performed the spleen cell sorting by flow cytometry to enrich the sample in ASCs and a targeted cell fusion to favor the production of hybridomas-secreting specific and affine antibodies.

## Results

### Characterization of ASCs phenotype using spleen cells from naive and immunized mice

To improve the efficiency of hybridoma generation from spleen cells, we aimed to select the most suitable ASCs, including plasmablasts and plasma cells, before cell fusion.

We selected a panel of five cell-surface markers specific to the B cell lineage (i.e., CD3, TACI, CD138, MHC class-II, and B220), and used this panel to identify ASCs among spleen cells from a naive and an immunized Balb-c mouse by flow cytometry (Fig. 1). We chose these markers since recent studies have underlined the B cell maturation process and have characterized mouse splenic plasma cells and plasmablasts using specific cell-surface proteins (11,22–24). We implemented a gating strategy (used for all experiments) to select single viable B cells from the spleen and determined the best marker combination to discriminate distinct ASC subsets. We first analyzed the expression of TACI and CD138 markers to select TACI^high^CD138^high^ plasmablasts (PB) and plasma cells (PC), as previously described (22). We observed that the immunization process often resulted in an increase of this spleen cell subset compared to the naive mice (i.e., 0.74% in Fig.1A vs 2.6% in Fig. 1B). Then, we examined the expression of MHC class II and B220, two well-known B cell markers whose expression decreases during the maturation of PB into PC (23,25,26). This gating strategy allowed us to distinguish three TACI^high^CD138^high^ cell populations according to B220 and MHC-II expression levels (Fig. 1), namely B220^high^MHC-II^high^ (P1), B220^low/int^MHC-II^high^ (P2), and B220^low^MHC-II^low/int^ (P3). In this experiment using NheB as the target antigen, P1 cells were more prominent in the naive mouse (about 66% in Fig.1A) than in the immunized mouse (about 24% in Fig.1B). P2 cells were underrepresented in the naive mouse (13% in Fig.1A) compared to the immunized mouse (54% in Fig.1B). P3 cells were present at similar levels, i.e., around 19%, for both naive and immunized mice.

**Figure 1:**
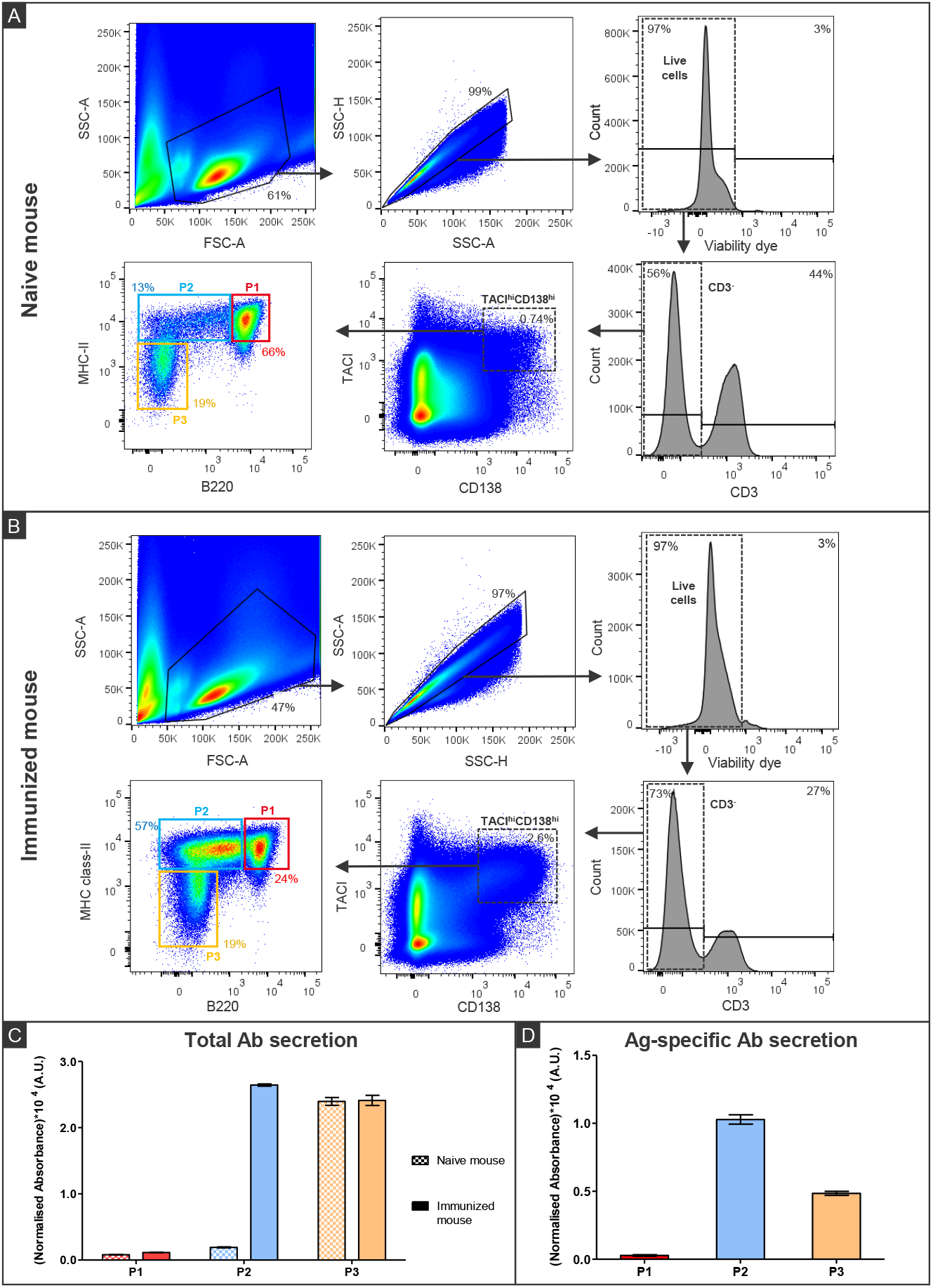
Analysis of ASCs from naive mouse and immunized mouse spleens. Flow cytometry analysis of ASCs from naive (A) vs. immunized (B) mice using the following gating strategy. A first gate was set on single lymphocytes through physical parameters (FSC-A vs. SSC-A) and on (SSC-H vs. SSC-A) to eliminate doublets, and a second gate on fluorescence parameters (viability dye) to exclude dead cells. Then, we focused on CD3^-^ cells to exclude the CD3^+^ T lymphocytes and targeted the CD138^high^TACI^high^ population that we subdivided into P1, P2 and P3 subsets according MHC-II and B220 expression. P1 cells are included in the red gate, P2 cells in the blue gate, and P3 cells in the orange gate. hi=high, int=intermediate, lo=low. Levels of total (C) and Ag-specific secreted antibodies (D) in cell culture supernatants 4 days after sorting of spleen cells from immunised and naive mice. For each cell population, the absorbance was normalized to the number of cells per well. Error bars = ± SD (duplicates). Data from a single experiment using two individual mice, one mouse having been immunized against NheB. Alt text: A) Four density plots and two distribution histograms characterizing size, granularity and five surface markers expression of spleen cells from a naive mouse revealing three cell populations. B) Similar plots and histograms as in panel A, using spleen cells from an immunized mouse, also revealing three cell populations. C) Bar chart comparing total antibody secretion levels from the three isolated cell populations shown in panels A and B. Three out of the six populations secrete antibodies. D) Bar chart comparing antigen-specific antibody secretion levels of the three isolated cell populations from immunized mice, revealing that two out of the three populations secrete antibodies.

We evaluated the antibody secretion levels in culture supernatant from each cell subset using ELISA assays to characterize the three identified cell populations. Specifically, we sorted around 10,000 cells from P1, P2, and P3 populations using spleen cells from the mice described above, and cultured them for four days in complete media supplemented with IL-4 and CD40L to mimic T-cell interactions and promote cell survival. We then measured the absorbance levels resulting from total secreted antibodies in cell supernatants, including IgG and IgM, by spectrophotometry (Fig. 1C). The absorbance levels resulting from Ag-specific secreted antibodies were only measured for the immunized mouse (Fig. 1D). For the naive mouse, P1 and P2 cells did not secrete antibodies at measurable levels, contrary to P3 cells whose secretion level was considerably higher. For the NheB-immunized mouse, P1 cells did not secrete any measurable antibodies (total or Ag-specific Fig.1C, D). In contrast, P2 and P3 cells secreted antibodies at high levels similar to P3 in naive mouse, with P2 secreting about twice as many Ag-specific antibodies than P3 cells (Fig.1D). P2 and P3 were thus consistent with an ASC phenotype, while P1 cells appeared as non-secreting for all tested conditions.

Additional data obtained using two other mice immunized with VIM-I showed similar results for phenotyping and antibody secretion profiles (Supplementary data Fig. S1 and Fig. S2), indicating that P1, P2, and P3 cells could also be found in splenocytes obtained using different sets of immunization conditions and were not specific to a single antigen or mouse.

### Evaluation of IgG expression and Ab secretion levels in ASCs

To characterize further the different P1, P2, and P3 cells, we used the membrane expression of IgG as an additional cell-surface marker and measured the secretion levels of the resulting cell subpopulations (Fig. 2). We generated these data from the same immunized mouse as in Fig.1. Results showed that 75% of P1 cells highly express membrane IgG, while 47% of P2 cells express membrane IgG at a medium level, and 28% of P3 cells at a medium to low level (Fig.2A). Following subsequent cell sorting of P1, P2, and P3 based on IgG expression, we performed dedicated immunoassays to measure Ab secretion levels in cell culture supernatants (Fig. 2B and Fig. 2C). Both IgG^+^ and IgG^−^ cells within the P2 and P3 cell subsets secreted antibodies at similarly high levels, although secretion levels from IgG^−^ cells tended to be slightly higher (Fig.2B). Regarding Ag-specific antibody secretion, once more IgG^+^ and IgG^−^ cells from the P2 and P3 cell subsets secreted antibodies (Fig.2C). Follow-up experiments conducted using spleen cells from mice immunized with VIM-I resulted in similar tendencies regarding IgG expression and secretion (supplementary data, Fig.S1 and Fig.S2). In these additional studies, we also evaluated IgM expression in P1, P2, and P3 cell subsets (supplementary data, Fig.S1A) and observed that P3, which only expressed membrane IgG at 24%, expressed IgM at 75%. Overall, these results showed that P2 and P3 can be considered as ASCs and it is interesting to note that they can both secrete and express Abs on the cell surface.

**Figure 2:**
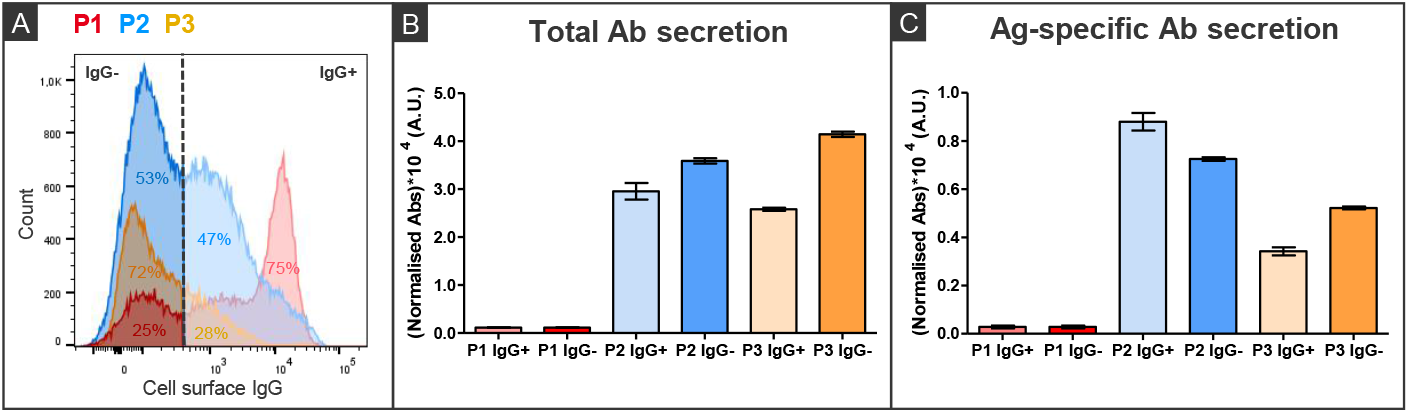
Analysis of membrane-expressed IgG and secreted Abs from P1, P2, and P3 ASCs from the spleen of a mouse immunized with NheB. ASCs expressing IgG at the cell surface were defined using IgG1, 2ab-PE-Vio770 antibody (A). The percentage of each ASC population expressing IgG (IgG^+^) or not (IgG^-^) is indicated in each histogram whose color is the same as the ASCs identified on the dot plots from Figure 1B. Around 5,000 cells of each ASCs subset were isolated. Detections of corresponding total antibodies (B) and Ag-specific antibody secretion (C) were performed in cell culture supernatants 4 days after cell sorting. For each cell population, the absorbance is normalized to the number of cells per well. Error bars = ± SD (technical duplicates). Representative data from a single experiment are shown. Additional experiments performed using two other mice immunised with VIM-I, gave similar results (See Supplementary data, Fig. S1 & S2). Alt text: A) Distribution histogram representing the cell surface IgG expression levels of the three cell populations identified in Fig. 1B. B) Bar chart comparing total antibody secretion levels from six cell sub-populations isolated from the three previously defined populations, based on either positive or negative IgG membrane expression. Four out of six sub-populations secrete antibodies. C) Bar chart comparing antigen-specific antibody secretion levels from the six isolated cell sub-populations. The same four sub-populations as in panel B secrete antibodies.

### Determination of the phenotype of hybridomas secreting Ag-specific Abs

We aimed at identifying more precisely which of the spleen cell populations has a higher probability of leading to hybridomas that can secrete Ag-specific Abs. Here, spleen cells from a mouse immunized against VIM were fused with myeloma cells using a conventional PEG-mediated cell fusion procedure. After ten days of culture in the selective medium, we analyzed, before cloning, a crude mixture of fused cells stained with the five-marker panel described above by flow cytometry. We also evaluated the expression level and specificity of membrane IgGs using biotinylated Ag combined with Streptavidin-PE as described by Parks et al. (27). The CD138 marker was not analyzed here since we considered all fused cells to be CD138^high^, due to its high expression on myeloma cells (supplementary data, Fig. S3). As expected, only 8% of cells were alive (FACS dot plots in Fig. 3A). Indeed, the majority of events on this plot were consistent with the presence of cell debris resulting from the death of non-fused cells due to the selective medium and the rarity of cell fusion events. Among the selected living cells, 87% of hybridomas were TACI^+^, of which 85% were MHC-II^+^B220^−^, thus corresponding to the identified ASC phenotype.

**Figure 3:**
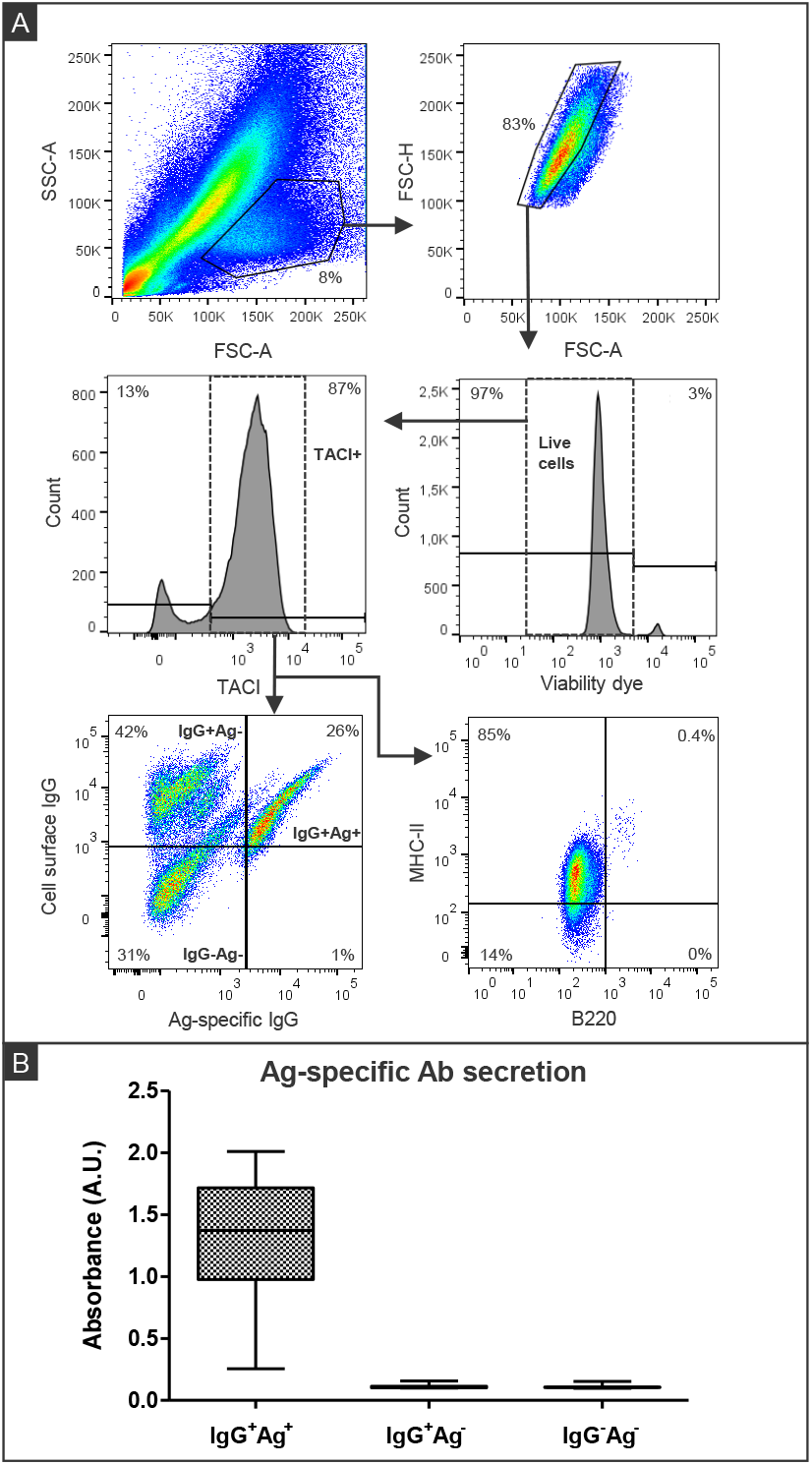
Identification of hybridoma secreting Ag-specific antibodies after fusion of spleen and myeloma cells. After ten days of culture in selective media, all hybridoma cells were collected and stained using a viability dye (Viobility dye 405/520), TACI-APC, B220-VioBlue, and MHC-II-PerCP-Vio700, IgG-PE-Vio770 antibodies and Ag-biotin/Streptavidin-PE. A first gate was set on viable hybridoma cell singlets through physical parameters (FSC-A vs. SSC-A) and on (FSC-H vs. FSC-A), and a second gate on fluorescent parameters (viability dye) to exclude dead cells. Then, we selected the TACI^+^ cells to analyse the expression of MHC-II and B220 together with cell surface IgG and their Ag specificity (A). Each of the 3 cell subsets identified (IgG^+^Ag^+^, IgG^+^Ag^-^, IgG^-^ Ag^-^) was isolated in single cell mode in separate wells. After 7 days, wells containing growing cells were tested for Ag specificity with dedicated immunoassays (B). Absorbance values are represented in box and whiskers plots with minimum, first quartile, median, third quartile and maximum values. Representative data from a single experiment using one mouse having been immunized against VIM-I is shown. An additional experiment performed using a second mouse, gave similar results (not shown). Alt text: A) Four density plots and two distribution histograms characterizing size, granularity and five markers expression of hybridoma cells. Three populations of hybridomas can be defined on the basis of the expression of antigen-specific or non-specific surface IgGs, B) Box plots comparing the antigen-specific antibody secretion levels from each of the three isolated cell populations identified in panel A. One out of the three populations secrete specific antibodies.

We then analyzed membrane expression of Ag-specific IgGs to divide TACI^+^ sorted cells into three new cell subsets: IgG^+^Ag^+^ (26%), IgG^+^Ag^−^ (42%), and IgG^−^Ag^−^ (31%). About four hundred cells from each subset were isolated using single-cell sorting mode by flow cytometry in separate 96-well plates. After 7 days of culture, we evaluated Ag-specific antibody secretion in the supernatants of the wells containing growing cells originated from a single isolated clone (35-43% of wells). Figure 3B shows that 100% of IgG^+^Ag^+^ cells secreted Ag-specific antibodies with a median absorbance value of 1.4, while none of the IgG^+^Ag^−^ and IgG^−^Ag^−^ cells secreted them at measurable levels. These results suggest that the only hybridoma cells capable of secreting Ag-specific antibodies also expressed them on the cell surface.

### Targeted fusion of TACI^high^CD138^high^ sorted spleen cells

To verify that fusion between ASCs and myeloma cells favored the generation of hybridomas secreting Ag-specific antibodies, compared to random spleen cells, we sorted TACI^high^CD138^high^ cells (i.e., expecting >60% of ASCs) from the spleen of a VIM-I-immunized mouse by flow cytometry. We achieved to isolate around 300 000 cells that we fused with myeloma cells using an electrofusion protocol. As a control, 300 000 non-sorted spleen cells from the same immunized mouse were fused in parallel (see workflow Fig. 4A). The fusion yield for both cell fusions was calculated using the number of wells containing growing hybridomas over the total number of plated wells (seven wells for each condition). Results showed 100% fusion efficiency for TACI^high^CD138^high^ cells, while for non-sorted spleen cells, cell growth was only observed in three wells (about 40% fusion yield).

**Figure 4:**
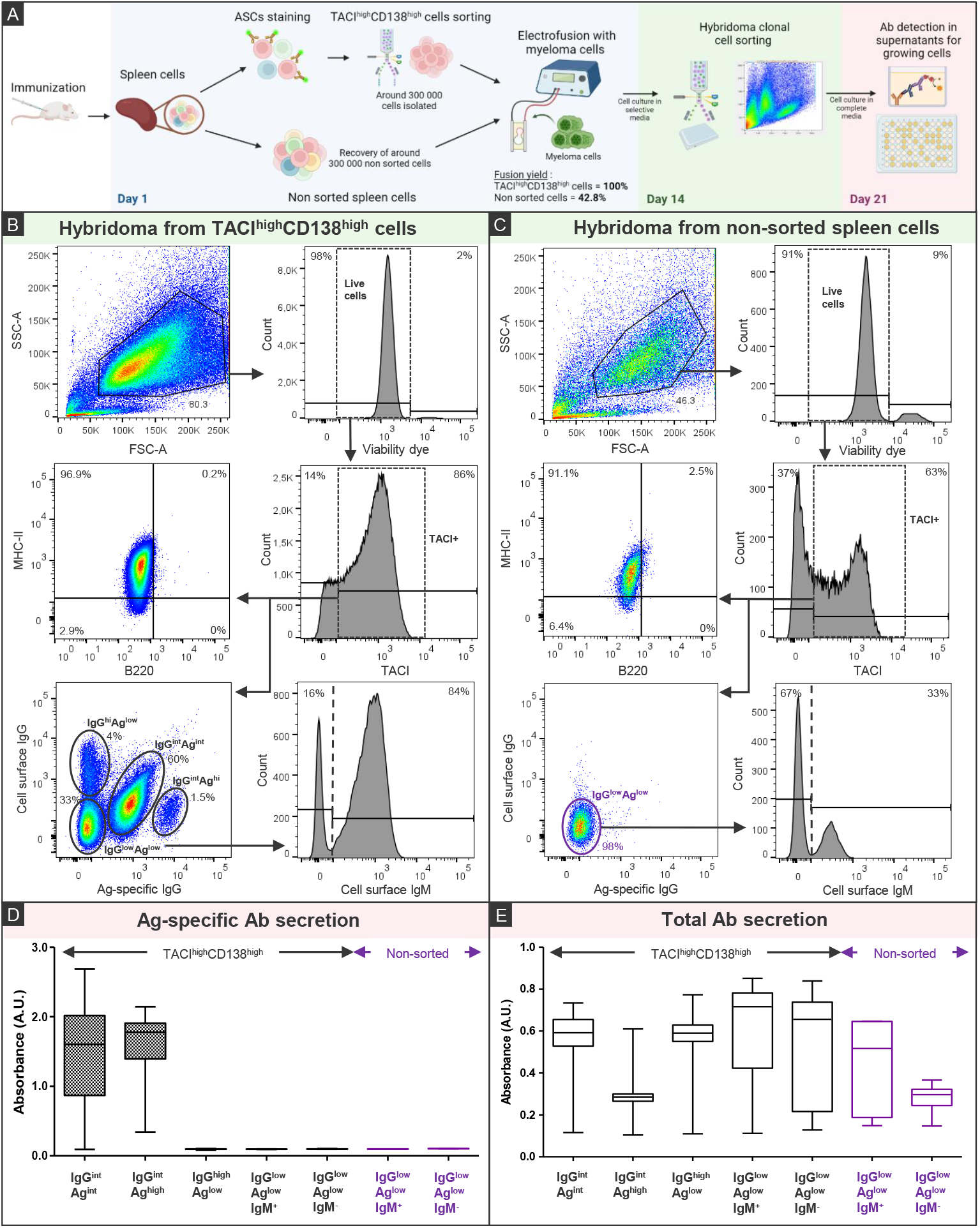
Identification of hybridoma secreting Ag-specific antibodies after targeted fusion of TACI^high^CD138^high^ spleen cells. Workflow of the entire process (A): TACI^high^CD138^high^ cells from a mouse immunized against VIM were sorted by FACS and electrofused with NS1 myeloma cells. As a control, non-sorted spleen cells from the same mouse were electrofused with NS1 myeloma cells. After fourteen days of culture in selective media, cells were collected and stained using TACI-APC, B220-VioBlue, MHC-II-PerCP-Vio700, IgG-PE-Vio770, IgM-APC-Vio770 antibodies and Ag-biotin/Streptavidin-PE. For both TACI^high^CD138^high^ fused cells (B) and non-sorted fused cells (C), a first gate was set on single viable hybridoma cells through physical parameters (FSC-A vs. SSC-A) and a second gate on fluorescent parameters (viability dye) to exclude dead cells. Then, we excluded the TACl^-^ cells to analyse the expression of MHC-II and B220 together with cell surface IgG and their Ag specificity. Each of the four cell subsets for TACI^high^CD138^high^ fused cells and the one subset for non-sorted fused cells were isolated in single cell mode in separate wells. For IgG^low^Ag^low^ cells, cell surface IgM expression was evaluated and IgM^+^ and IgM^-^ cells were also isolated in single cell mode in separate wells. After 7 days, wells containing growing cells were tested for Ag specificity (D) and total Ab (E) secretion with dedicated immunoassays. Absorbance values are represented in box and whiskers plots with minimum, first quartile, median, third quartile and maximum values. An additional experiment performed using another mouse, gave preliminary results (See Supplementary data, Fig. S6). high=hi/intermediate=int/low=lo. Alt text: A) Drawing of the workflow of the targeted electrofusion process. B) Three density plots and three distribution histograms characterizing size, granularity and six markers expression of hybridomas obtained after fusion of antibody secreting cells, revealing five populations on the basis of positive or negating surface expression of IgG, antigen-specific IgG and IgM. C) Similar plots and histograms as in panel B, but using hybridomas obtained after fusion with random spleen cells, revealing only two populations based on the same surface markers. D) Box plots comparing antigen-specific antibody secretion levels from the seven isolated cell populations described in panels B and C. Two out of seven cell populations secrete antibodies. E) Box plots comparing total antibody secretion levels of the same seven populations. All cell populations secrete antibodies at varying levels.

After 14 days of incubation in the selective medium, we analyzed the membrane expression of TACI, B220, MHC-II, IgG, and the Ag specificity of the resulting hybridomas from both tested conditions by flow cytometry (Figs. 4B and 4C). After gating for TACI^+^, more than 90% of selected cells displayed an ASC phenotype, with MHC-II^+^B220^−^ expression, for both cell fusions. Subsequently, the analysis of cell-surface expressed Ag-specific IgGs revealed distinct profiles depending on the initial cells used for fusion. For the hybridomas produced by fusion with non-sorted cells, there was only one population with IgG^low^Ag^low^ phenotype. For the hybridomas produced by fusion with TACI^high^CD138^high^ sorted cells, we could distinguish four different cell subsets: the major one was IgG^int^Ag^int^ (60% of cells), a second only present at 1.5% was IgG^int^Ag^high^, and the two other were IgG^high^Ag^low^ and IgG^low^Ag^low^ found at 4% and 33%, respectively. IgM expression analysis on IgG^low^ cells from both non-sorted and TACI^high^CD138^high^ fused cells showed that these respective subsets expressed IgM at 33% and 84%.

We then clonally isolated the following cell subsets: IgG^int^Ag^int^, IgG^int^Ag^high^, IgG^high^Ag^low^, IgG^low^Ag^low^IgM^+^, and IgG^low^Ag^low^IgM^−^ from both cell fusions and evaluated Ag-specific Ab secretion levels in supernatants after 7 days of culture (Fig. 4D). We observed that only cells expressing Ag-specific IgGs at the cell surface could secrete them (median values of 1.59 and 1.78 for IgG^int^Ag^int^ and IgG^int^Ag^high^, respectively). It is important to note that no hybridomas were able to secrete Ag-specific Abs for the non-sorted fused cells.

Since the other cell subsets displayed an ASCs phenotype without secretion of Ag-specific Abs, we investigated the secretion of total Abs (IgG and IgM) in cell supernatants (Fig. 4E). Interestingly, we found that all cell subsets secreted Abs at varying levels. While IgG^int^Ag^int^, IgG^high^Ag^low^, IgG^low^Ag^low^IgM^+^, and IgG^low^Ag^low^IgM^−^ from TACI^high^CD138^high^ fused cells secreted high levels of total Abs (median value > 0.5), IgG^int^Ag^high^ from sorted cells and IgG^low^Ag^low^IgM^+^, IgG^low^Ag^low^IgM^−^ from non-sorted cells secreted lower levels of Abs (median value ≤ 0.5). Isotyping was then conducted for each subset to ascertain the phenotype between IgG and IgM (see supplementary data, Fig. S4). We observed that IgM^+^ cells secreted IgM antibodies, while IgG^int/high^ cells secreted IgG1_kappa_ antibodies, and IgG^int^Ag^high^ and IgG^low^Ag^low^IgM^−^ from non-sorted cells secreted IgA antibodies.

## Discussion

The present study provides a better understanding and an improvement of the hybridoma technology by identifying and characterizing antibody-secreting cells (ASCs) as the most suitable partners among spleen cells for the generation of hybridomas secreting Ag-specific antibodies.

ASCs are differentiated from activated B cells following immunization. We analyzed several combinations of markers specific to the B cell lineage to select this population over other spleen cells. The cell-surface markers were chosen due to their upregulation or downregulation during B cell maturation. A panel including CD3, TACI, CD138, MHC-II, and B220 markers allowed the definition of three B cell subsets in naive and immunized mice spleen, named P1, P2, and P3. In this study, the P1 cell subset TACI^high^CD138^high^B220^high^MHC-II^high^ appeared as non-secretory cells for both tested conditions but displayed a high cell surface expression of IgG for the immunized mouse. A P1* cell subset from VIM1-immunized mouse (not stained for TACI, Fig. S5) showed that these surface IgGs were specific for the target antigen. Therefore, although it did not secrete Abs, this cell subset is similar to what is usually called early plasmablasts (28). The P2 cell subset TACI^high^CD138^high^B220^low/int^MHC-II^high^ are ASCs along with the P3 cell subset TACI^high^CD138^high^B220^low^MHC-II^low/int^ since both expressed and secreted IgG, as previously described (9,29–31). However, while P2 and P3 cells expressed membrane IgGs, P3 cells expressed mainly IgM. In addition, both cell subsets secreted Ag-specific Abs (IgG+IgM), but at a lower proportion over total Abs for P3 cells. These results suggest that P3 cells could secrete low-affinity Ag-specific Abs.

Comparison of the representation of the three cell subsets in the immunized and naive mice revealed a significant decrease of P1 cells under immunization and a substantial increase of P2 cells in the spleen, suggesting a maturation process of P1 cells into P2 cells through Ag exposure. On the other hand, P3 cells were present in the same proportion in both naive and immunized mice. Taking into account the phenotype, IgM membrane expression, and Ab secretion levels of P3 cells, we suggest that these plasma-like cells may originate from an extrafollicular response in the spleen that does not involve T cell interaction, which could lead to the development of plasma cells that secrete low-affinity IgM (10,32,33). P1 and P2 cells, however, seem to derive from a germinal center response where T cell interactions allow B cells class switch and affinity maturation by somatic hypermutations that result in the formation of highly Ag-specific cells. Nevertheless, P2 cells, which conserve PB characteristics with high MHC-II expression, surprisingly secreted high levels of Ag-specific antibodies. This observation could be explained by the fact that P2 cells underwent further maturation in plate wells after cell sorting due to IL-4 and CD40L enrichment in the culture media, leading to PC formation. However, our additional experiments demonstrated no significant influence of IL-4 and CD40L, neither of the incubation time following cell sorting on Ab secretion. We also expected to identify a higher percentage of Ag-specific IgG-secreting spleen plasma cells with P3 cell phenotype, but maybe some of these cells have already left the spleen and migrated to plasma cell niches in the bone marrow (34).

Therefore, P2 cells appeared as the most interesting ASCs to fuse and generate Ag-specific IgG-secreting hybridomas. To confirm this hypothesis, we analyzed the phenotype of a hybridoma mixture obtained after cell fusion using our five-marker panel, and it revealed that these cells had an ASC-like phenotype with high expression of TACI and CD138, a low expression on B220 and an intermediate/high expression of MHC class II. Moreover, ELISA analysis demonstrated that all clones secreting Ag-specific Abs also expressed Ag-specific IgGs at the cell surface. We can thus hypothesize that this IgG expression in hybridomas may originate from spleen cells, probably from P2 and P3 ASCs. Furthermore, the fact that among TACI^+^ hybridoma cells, 85% were MHC-II^+^B220^−^ (like P2 and P3 cells) seems to confirm that fusion events occur preferentially with ASCs. The almost exclusive ASC phenotype of hybridoma cells could be explained by various factors influencing the cell fusion process, including the cell type, as well as cell size and the curvature of the cell membrane (either for PEG-based (35–38) or electrofusion (39–41)). We speculate that there is a higher probability for NS1 myeloma cells, which are abnormal PC, to undergo fusion with cells of a similar type and a comparable size. Due to their extended Golgi apparatus, ASCs are larger in size and could be a more suitable fusion partner for NS1 cells than other B cells. Notably, the ASC phenotype of hybridomas can not be attributed to NS1 cells that are TACI^−^, MHC-II^−^, and B220^−^ (supplementary data, Fig.S3). Additional investigations focusing on the fusion of myeloma cell lines different from NS1 and using a large panel of target antigens may provide further insight into this question.

Finally, we demonstrated that sorting TACI^high^CD138^high^ cells to increase the proportion of ASCs before fusion, successfully enhanced both the fusion yield and the number of generated Ag-specific-secreting hybridomas (Fig. 4 and supplementary data, Fig.S6). In contrast, the fusion of non-sorted spleen cells (i.e., when starting with 300,000 cells) did not produce any Ag-specific-secreting hybridomas. These results could be due to the relatively low percentage of ASCs in the spleen (0.5-2%), which reduces the probability of successful fusion events, especially when dealing with a limited number of cells. Enriching the spleen cell suspension with ASCs allowed the generation of two types of hybridoma cells: IgG-expressing and IgG-secreting hybridomas, some of the IgG being Ag-specific mAbs, and the IgM-expressing and IgM-secreting hybridomas, expected to display low affinity for the target antigen. By linking these results with spleen cell profiles, we can assume that IgG-producing cells may derive from P2 cells, while IgM-producing hybridomas may originate from P3 cells. Although the results from PEG-based fusion using a high number of non-sorted spleen cells or from electrofusion using a small number of ASC-enriched cells cannot be directly compared, the last method led to a five-fold increase in the percentage of generated Ag-specific secreting-hybridoma cells. This estimation was based on the IgG^+^Ag^+^ cells percentage reported to the total number of viable hybridomas counted before cell sorting. In addition, working with fewer cells simplifies the post-fusion process by reducing plate handling and facilitating downstream selection.

In conclusion, this study demonstrated that selecting ASC populations before cell fusion significantly enhances the efficiency of hybridoma production. Our findings suggest that fusion between spleen and myeloma cells primarily occurs with ASCs, which might explain the low frequency of fusion events in the conventional method and the improvements of our approach. In addition, by isolating ASC populations first, we were able to streamline the hybridoma technology procedure.

Overall, it is evident that the enhancement of the hybridoma technology will facilitate a more wide spread deployment of this technique, with the potential to reduce the use of animal models and to produce human hybridomas by isolating and fusing ASCs directly from patients.

## Data limitations and perspectives

The FACS-based isolation of TACI^high^CD138^high^ cells remains labor-intensive and costly in terms of time and reagents. Efforts are underway to develop magnetic cell sorting as a more efficient alternative for ASC selection from spleen cells. Moreover, future studies will be focused on the efficient isolation and exclusive fusion of Ag-specific ASCs to generate only Ag-specific secreting-hybridomas.

## Materials and Methods

### Mice immunization

Balb-c mice (8 to12 weeks), bred in the animal care unit at CEA, were immunized intraperitoneally with 50 μg per injection of purified antigen with alum hydroxide (1:1) after being anesthetized with isoflurane delivered through a vaporizer. The mice received four immunizations every three weeks and were left at rest for two months after the last injection. Before use, the animals were injected intravenously with 50 µg of antigen for three successive days and then were killed by cervical dislocation.

The NheB-immunized mice were already available in our facilities and used in the first experiments. NheB (produced and supplied by ANSES) is a subunit derived from the enterotoxin Nhe, which is secreted by Bacillus cereus and is responsible for food poisoning. For the following experiments, mice were immunized against VIM-I, an enzyme responsible for antibiotic resistance.

### Cell culture

The myeloma cell line NS1 was cultured in RPMI 1640 supplemented with 10% heat-inactivated fetal calf serum (FCS), 2 mM glutamine, 2 mM sodium pyruvate, 1% Penicillin/Streptomycin, 1% MEM with non-essential amino acids (all purchased from Sigma Aldrich) at 37°C and 7% CO_2_.

### PEG-mediated cell fusion

Splenocytes from mice immunized with the target antigen were recovered. Splenocytes and myeloma cells were washed and then suspended in RPM1 1640 serum-free medium (Sigma) at 3:1 ratio. Then, both cells were centrifuged together at 1,000 rpm for 10 min. After removing the supernatant, one milliliter of 50% PEG 1500 (Sigma) was added dropwise to the cell pellet over 1 min with gentle stirring. After a 90 s incubation at 37°C, 50 ml of the serum-free culture medium were added dropwise. The cells were then centrifuged for 10 min at 1,000 rpm, washed twice, and resuspended in selective medium (RPM1 1640 supplemented with 15% heat-inactivated FCS, 1X hypoxanthine (ThermoFisher), 2 mM glutamine, 2 mM sodium pyruvate, 1% Penicillin/Streptomycin (all purchased from Sigma Aldrich), called after 15% FCS complete medium. Cells were finally distributed in 6-well microtiter plates containing macrophages as feeder layer (5×l0^3^ Balb/c peritoneal exsudate cells per well) and incubated at 37°C and 7% CO2.

### Cell electrofusion

Spleen cells were recovered from cell sorting and added to myeloma cells at 1:1 ratio. Cells were then washed with RPMI 1640, suspended in 80µL of electrofusion buffer (100mM Sorbitol, 0.5mM MgCl_2_, 0.1mM Ca (CH_3_COO)_2_, 1mg/mL BSA, pH=7.5), transferred into 0.1cm gap electroporation cuvette (BIORAD) and centrifuged for 5 min at 300 xg. The cuvette was placed in the Gene Pulser Xcell Electroporation System (BIORAD) and electrofusion was performed using a Square Wave Protocol of 1500 kV/cm electric field, 50 µs pulse duration, 3 repetitions with 0.1s interval. After electrofusion, the cuvette was incubated for 10 min at 37°C. Cells were then slowly suspended, transferred into a 50 mL tube and diluted at about 1×10^5^ cells/mL with selective medium. Finally, after 20 min incubation at 37°C, cells were distributed in 96-well plates and incubated at 37°C and 7% CO2.

### ASCs cell sorting

Mouse spleens were harvested and crushed through a 40µm cell strainer (ThermoFisher) to extract splenocytes. Red bloods cells were lysed using Red Blood Cell Lysis Buffer (Sigma). Cells were then washed twice with PBS (−)CaCl_2_ (−)MgCl_2_ (Sigma) and stained for 15 min at 4°C according to manufacturer instructions using the following fluorescent antibodies (Miltenyi Biotec): Viobility 405/520 Fixable Dye, CD3e-Vio^®^Bright FITC, CD267(TACI)-APC, CD138-PE, CD45R(B220)-Vio^®^Blue, MHC Class II-PerCP-Vio^®^700, IgG2ab-PE-Vio^®^770, IgG1-PE-Vio^®^770.Then, cells were washed twice with PBS and sorted using BD FACSAria™ III and BD FACSDiva software. Sorted cells were recovered in RPMI 1640 supplemented with 15% heat-inactivated FCS, 2 mM glutamine, 2 mM sodium pyruvate, 1% Penicillin/Streptomycin, 1% MEM with non-essential amino acids, 2% HFCS (Hybridoma Fusion and Cloning Supplement) (all from Sigma) and incubated at 37°C and 7<u>%</u> CO_2_ for several days. FACS data analysis was performed using the FlowJo software V10.10.0 (BD Biosciences).

### Hybridoma cell sorting

Cells were washed twice with PBS (−)CaCl_2_, (−)MgCl_2_ (Sigma) and incubated for 30 min at 4°C with biotinylated antigen at 5µg/mL. Then, cells were washed and stained for 15 min at 4°C according manufacturer instructions using fluorescent antibodies (Miltenyi Biotec): Viobility 405/520 Fixable Dye, CD3e-Vio^®^Bright FITC, CD267(TACI)-APC, CD138-PE, CD45R(B220)-Vio^®^Blue, MHC Class II – PerCP-Vio^®^700, IgG2ab-PE-Vio^®^770, IgG1-PE-Vio^®^770 and Streptavidin-PE. After staining, cells were washed twice with PBS and sorted using BD FACSMelody™. Sorted cells were recovered in 15% FCS complete medium and incubated in 37°C and 7% CO_2_ for several days. FACS data analysis was performed using FlowJo software V10.10.0.

### Antibody detection in cell culture supernatants

100µL of cell culture supernatants were transferred into microtiter plates coated with goat anti-mouse IgG+IgM antibodies (Jackson Immunoresearch laboratories) and incubated overnight at 4°C. For Ag-specific antibodies detection, plates were washed and incubated with 100µL/well of biotinylated antigen at 100ng/mL for 4h at room temperature (RT). AcetylCholinesterase (AChE)-conjugated streptavidin was then added at 1 Ellman units [UE]/mL after washing and incubated for 1h at RT. For total antibodies detection, plates were washed and incubated with 100µL of goat anti-mouse-AChE antibodies at 2 Ellman units [UE]/mL for 1h at RT. Finally, plates were washed and incubated with 200 µL of Ellman’s reagent (protected from light) and the absorbance was measured at λ = 414 nm after 30min or 1h using Epoch spectrophotometer (BioTek instruments).

### Antibody isotyping

Antibody isotyping was performed using Pierce Rapid ELISA Mouse mAB Isotyping Kit from Invitrogen and according fabricant’s instructions. Briefly, cell culture supernatants were diluted at 1:50 in 1 mL Tris Buffered Saline. Then, 50 µL of the diluted supernatant and 50 µL of Goat anti-Mouse IgG+IgA+IgM-HRP were added to each well of the 8-well pre-coated strip, and covered for 1h at RT. After three washings using the wash buffer (potassium phosphate buffer 10mM, 0.05% Tween 20), 75µL of TMB substrate was added to each well and incubated for 15min protected from light. Then, 75µL of Stop Solution was added and absorbance values were measured at λ = 450 nm using Epoch spectrophotometer (BioTek instruments).

## Abbreviations

mAb: monoclonal antibody
Ab: Antibody
ASC: Antibody-Secreting Cell
FACS: Fluorescence-Activated Cell Sorting
Ig: Immunoglobulin
Ag: Antigen
PEG: Polyethylene glycol
PB: Plasmablast
PC: Plasma cell
ELISA: Enzyme-Linked ImmunoSorbent Assay

## Data Availability Statement

This study includes no data deposited in external repositories

## Ethics Statement

All experiments were performed in compliance with French and European regulations on the care of laboratory animals (European Community Directive 86/609, French Law 2001-486, 6 June 2001) and with the agreements of the Ethics Committee of the Commissariat à l’Energie Atomique (CEtEA ‘Comité d’Ethique en Expérimentation Animale’ n° 44) no. 12-026 and 15-055 delivered to S.S. by the French Veterinary Services and CEA agreement D-91-272-106 from the Veterinary Inspection Department of Essonne (France).

## Author Contributions

Fanny Rousseau designed and performed experiments, analyzed the data, and wrote the paper. Catherine Menier contributed to FACS experiments and data analysis and revised the manuscript. Patricia Brochard contributed to fusion experiments and provided technical guidance. Stéphanie Simon supervised the complete study. Anne Wijkhuisen and Karla Perez-Toralla contributed to some experiments, conceived the manuscript, wrote the introduction, and revised the manuscript. All authors read and approved the final manuscript.

## Acknowledgments

We thank Marc Plaisance, Claire Vagneux and Mequa Maatoug (CEA, France) for assistance with animal experiments. We thank Mathilde BONIS and Sophie LIUU (Anses, France) for NheB toxin supply. Some Figures were created using BioRender.com. This work was funded by CEA, France. We used Grammarly (free version 14.1217.0) occasionally to improve grammar, punctuation and sentence clarity on the final and revised manuscript. Each suggestion was reviewed and modified independently before adapting it.

