## Supporting information for "Targeted fusion of Antibody-Secreting Cells: unlocking monoclonal antibody production with hybridoma technology"

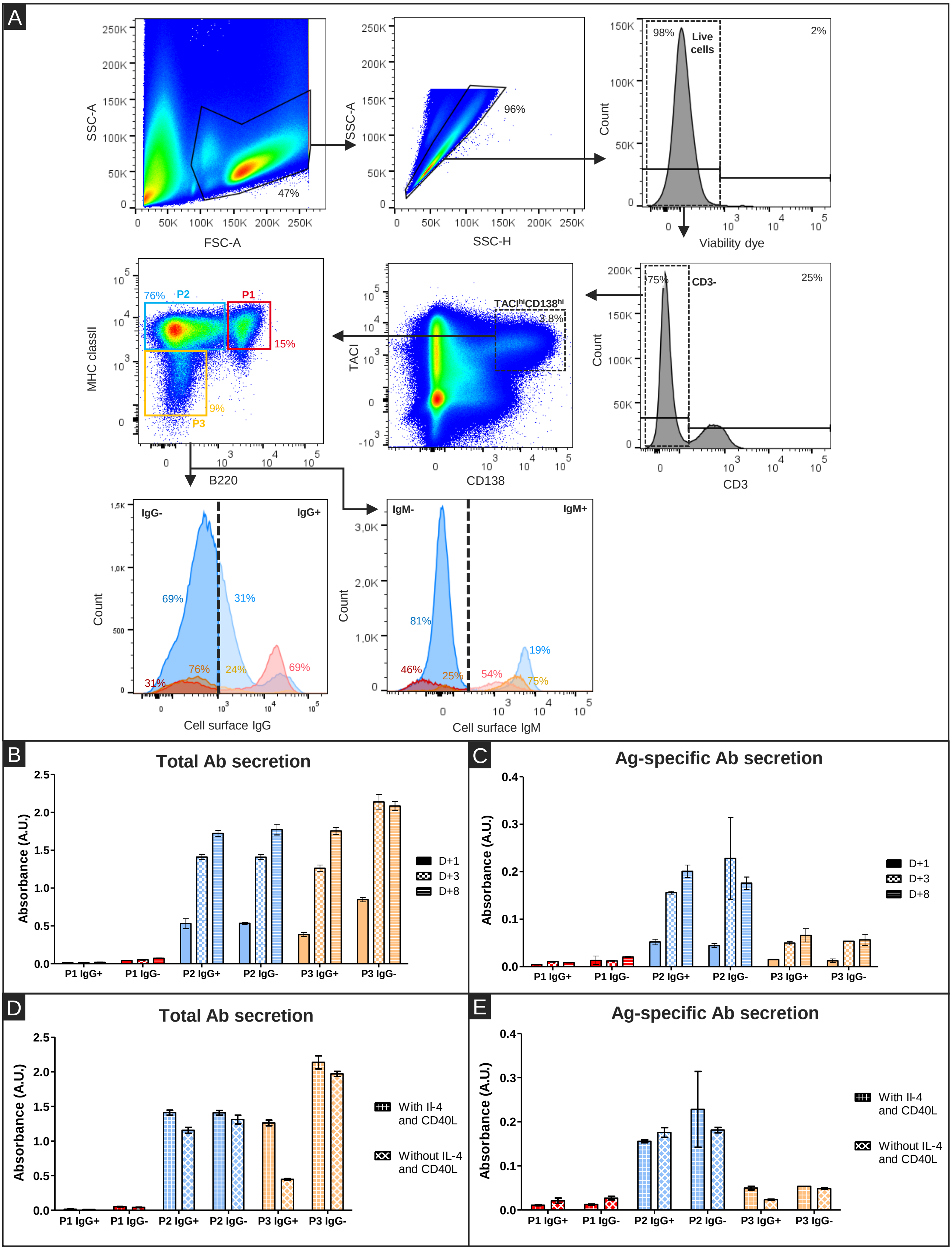


**Figure S1: Identification of antibody secreting cells (ASCs) and antibody expressing cells from a mouse immunized against VIM-I.**
ASCs were defined using CD3-FITC. TACI-APC. CD138-PE. B220-VioBlue. and MHC-II-PerCP-Vio700 antibodies by flow cytometry analysis using the following gating strategy. A first gate was set on single lymphocytes through physical parameters (FSC-A vs. SSC-A) and on (SSC-H vs. SSC-A) to eliminate doublets. and a second gate on fluorescence parameters to exclude dead cells. Then. we excluded the $CD3^{+}$ T lymphocytes and targeted the $CD138^{high}TACI^{high}$ population that we subdivided into P1. P2 and P3 subsets according MHC-II and B220 expression. P1 cells are included in the red gate. P3 cells in the orange gate and P2 cells in the blue gate. IgG expressing cells and IgM expressing cells were defined using IgG1.2ab PE-vio770 and IgM APC-vio770 antibodies and are represented with subsets corresponding colours. For $IgG^{+}$ and $IgG^{-}$ subset. around 5000 cells/subsets were isolated except for P1 $IgG^{-}$ and P3 $IgG^{+}$ with 3000 cells/subsets. ELISA immunoassays of total (B) and Ag-specific secreted antibodies (C) in cell culture supernatants 1. 3. 8 days after cell sorting. ELISA quantification of total (D) and Ag-specific secreted antibodies (E) in cell culture supernatants supplemented or not with CD40L and IL-4 three days after cell sorting. For each cell population. the absorbance of the total number of cells per well is represented. Error bars = ± SD (technical duplicates). Representative data from a single experiment using one mouse having been immunized against VIM-I.

**Figure S1:** Alt text

A) Four density plots and two distribution histograms characterizing size, granularity and five surface markers expression of spleen cells from an immunized mouse, revealing 3 populations. Two additional distribution histograms for cell surface IgG and IgM expression levels of the 3 identified populations. B) Bar chart comparing total antibody secretion levels at three different days from six cell sub-populations isolated from the three previously defined populations, based on either positive or negative IgG membrane expression. Four out of six sub-populations secrete antibodies. C) Similar charts as B for antigen-specific antibody secretion levels. The same four sub-populations as in panel B secrete antibodies. D) Bar chart comparing total antibody secretion levels in presence or in absence of IL-4 and CD40L from six cell sub-populations isolated from the three previously defined populations, based on either positive or negative IgG membrane expression. Four out of six sub-populations secrete antibodies. E) Similar charts as D for antigen-specific antibody secretion levels. The same four sub-populations as in panel D secrete antibodies.

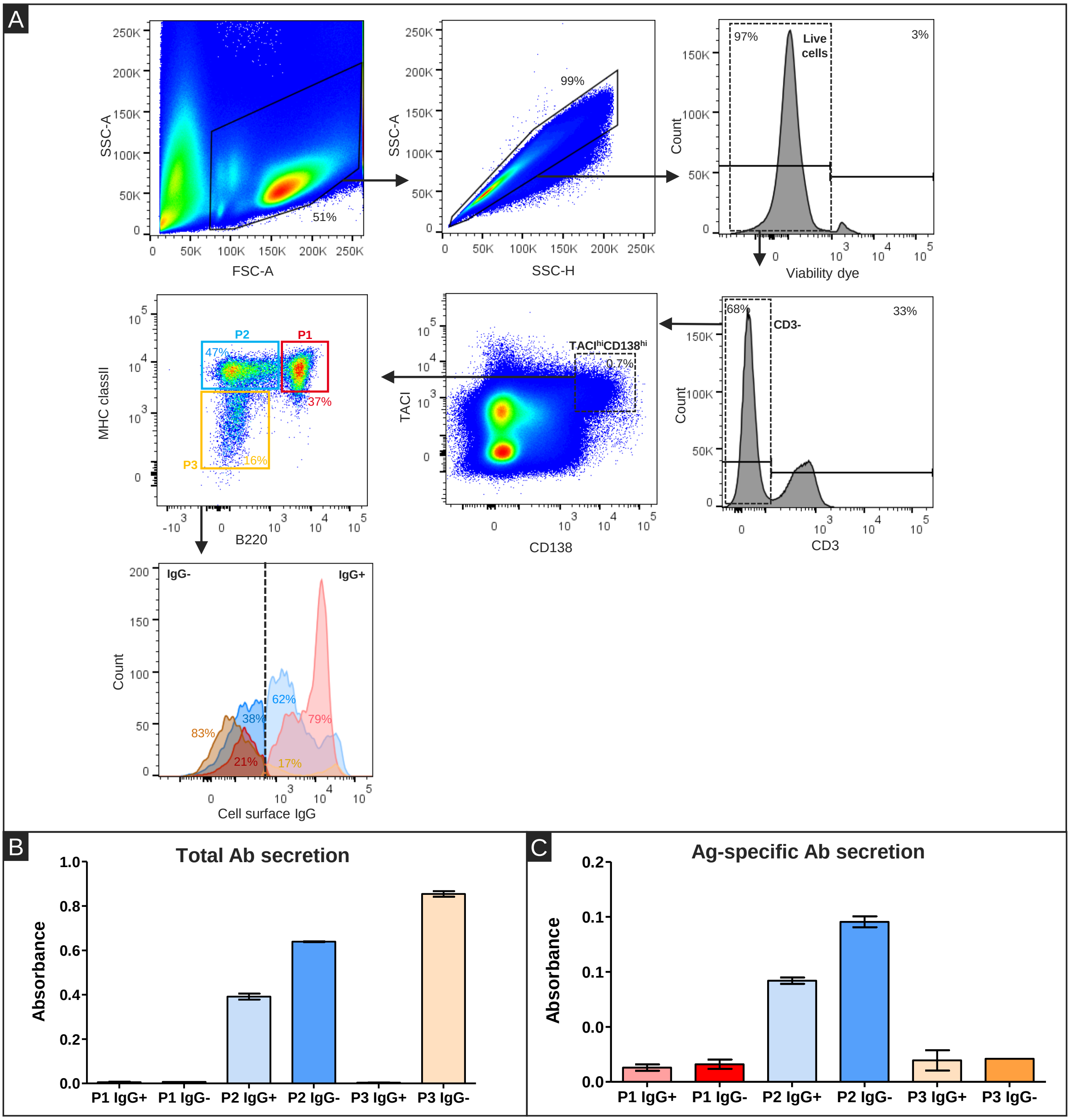


**Figure S2: Identification of antibody secreting cells (ASCs) and antibody expressing cells from a mouse immunized against VIM-I.**
ASCs were defined using CD3-FITC. TACI-APC. CD138-PE. B220-VioBlue. and MHC-II-PerCP-Vio700 antibodies by flow cytometry analysis using the following gating strategy. A first gate was set on single lymphocytes through physical parameters (FSC-A vs. SSC-A) and on (SSC-H vs. SSC-A) to eliminate doublets. and a second gate on fluorescence parameters to exclude dead cells. Then. we excluded the $CD3^{+}$ T lymphocytes and targeted the $CD138^{high}TACI^{high}$ population that we subdivided into P1. P2 and P3 subsets according MHC-II and B220 expression. P1 cells are included in the red gate. P3 cells in the orange gate and P2 cells in the blue gate. Around 5000 cells/subset were isolated except for P3 $IgG^{+}$ with less than 1000 cells isolated. ELISA detection of total (B) and Ag-specific secreted antibodies (C) in cell culture supernatants 4 days after cell sorting. For each cell population. the absorbance of the total number of cells per well is reprensented. Error bars = ± SD (technical duplicates). Representative data from a single experiment using one mouse having been immunized against VIM-I.

Alt text: A) Four density plots and two distribution histograms characterizing size, granularity and five surface markers expression of spleen cells from an immunized mouse, revealing 3 populations. One additional distribution histogram for cell surface IgG expression levels of the 3 identified populations. B) Bar chart comparing total antibody secretion levels from six cell sub-populations isolated from the three previously defined populations, based on either positive or negative IgG membrane expression. Three out of six cell populations secrete antibodies. C) Similar charts as B for antigen-specific antibody secretion levels. Two out of six cell populations secrete antibodies.

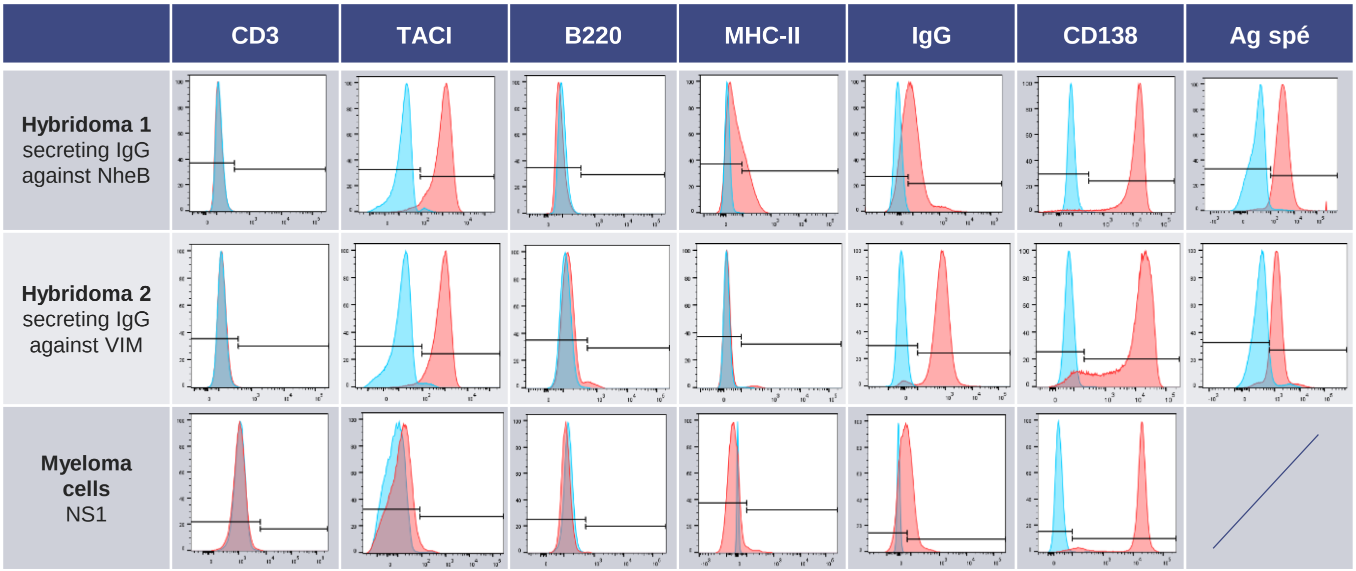


**<u>Figure S3</u>: Phenotype determination of hybridoma and NS1 cell lines**.
Expression of the five markers panel previously defined plus membrane Ag-specific Ab was evaluated by flow cytometry on two hybridoma cell lines (first and second rows) and the NS1 myeloma cell line (third row). Unstained cells were used as control (blue histograms). The MFI was calculated based on negative events (black gates). Data are represented normalized to the mode.

<u>Alt text:</u> Table showing distribution histograms of seven markers expression levels for three different cell lines. Each histogram shows a comparison between stained and unstained cells as control.

| | Hybridoma from TACI$^{high}$CD138$^{high}$ cells | | | | | Hybridoma from non-sorted cells | |
|---|---|---|---|---|---|---|---|
| | IgG$^{int}$Ag$^{int}$ | IgG$^{high}$Ag$^{low}$ | IgG$^{low}$Ag$^{low}$IgM- | IgG$^{low}$Ag$^{low}$IgM+ | IgG$^{int}$Ag$^{high}$ | IgG$^{low}$Ag$^{low}$IgM+ | IgG$^{low}$Ag$^{low}$IgM- |
| **IgG1** | 3.912 | 3.852 | >4 | 0.076 | 0.104 | 0.061 | 0.109 |
| **IgG2a** | 0.064 | 0.058 | 0.087 | 0.079 | 0.064 | 0.064 | 0.071 |
| **IgG2b** | 0.08 | 0.073 | 0.09 | 0.065 | 0.064 | 0.082 | 0.061 |
| **IgG3** | 0.073 | 0.112 | 0.075 | 0.105 | 0.092 | 0.083 | 0.085 |
| **IgA** | 0.062 | 0.059 | 0.058 | 0.058 | 0.767 | 0.055 | 0.17 |
| **IgM** | 0.059 | 0.057 | 0.056 | 2.613 | 0.073 | 0.179 | 0.057 |
| **Kappa** | 0.485 | 0.63 | 0.713 | 0.514 | 0.138 | 0.074 | 0.066 |
| **Lambda** | 0.059 | 0.061 | 0.057 | 0.059 | 0.061 | 0.054 | 0.054 |

**Figure S4: Isotyping of secreted Abs from sorted hybridoma cells.**
The supernatant of each subset identified in Figure 4 was tested using an ELISA Mouse mAb isotyping kit for all isotype (IgG1, IgG2a, IgG2b, IgG3, IgA and IgM). The absorbance values obtained are represented in the following table. Considered positive values (>0.150) are represented in green.

Alt text: Table showing the isotype data from seven different cell subsets identified in Figure 4 in the main text.

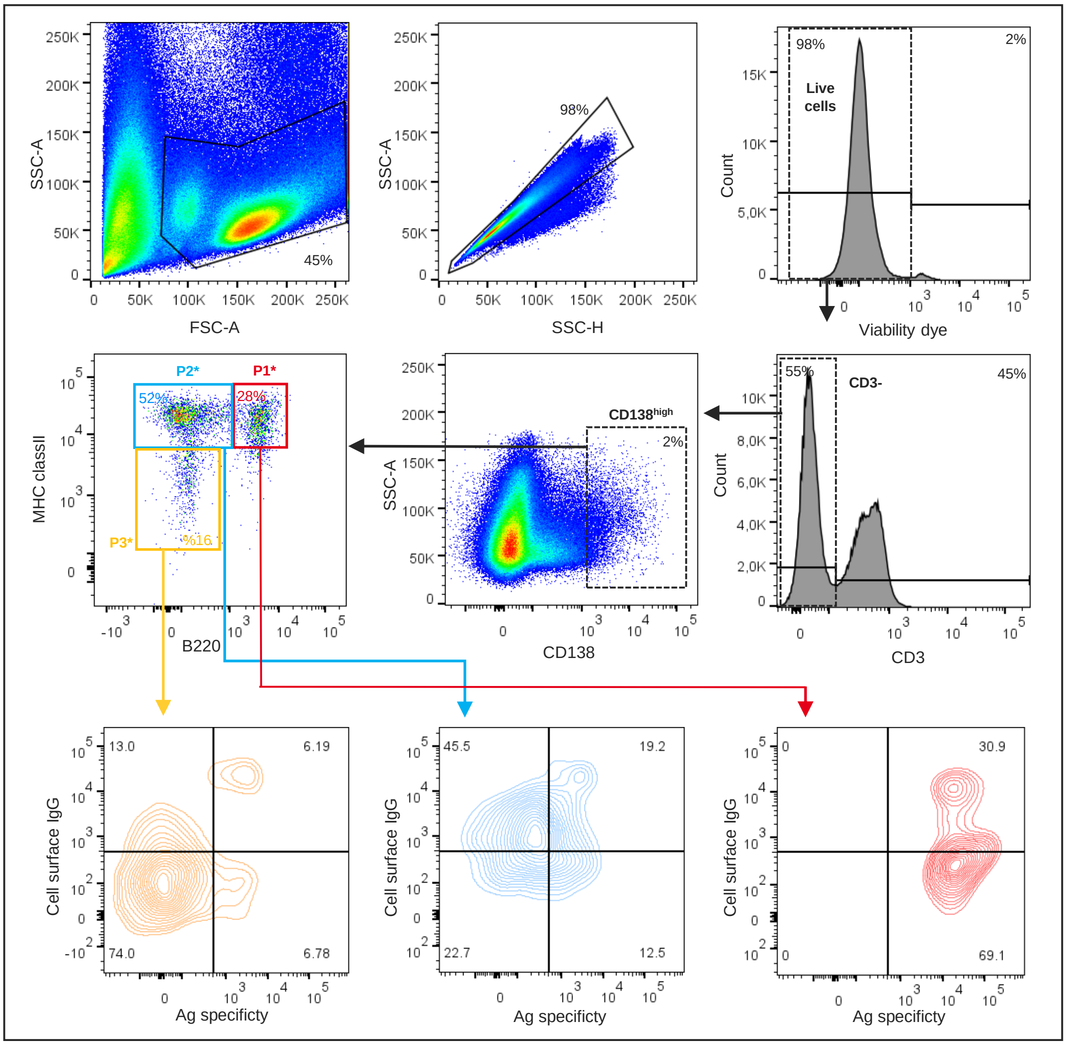


**<u>Figure S5:</u> Identification of Ag-specific IgG expression on antibody secreting cells (ASCs) from a mouse immunized against VIM-I.**

ASCs were defined using CD3-FITC. CD138-PE. B220-VioBlue. and MHC-II-PerCP-Vio700 antibodies by flow cytometry analysis using the following gating strategy. A first gate was set on single lymphocytes through physical parameters (FSC-A vs. SSC-A) and on (SSC-H vs. SSC-A) to eliminate doublets. and a second gate on fluorescence parameters to exclude dead cells. Then. we excluded the $CD3^{+}$ T lymphocytes and targeted the $CD138^{high}$ population that we subdivided into P1*. P2* and P3* subsets according MHC-II and B220 expression. P1* cells are included in the red gate. P2* cells in the blue gate and P3* cells in the orange gate. For each subset. Ag-specific IgG expression were defined using IgG1.2ab-PEvio770 antibodies combined with Ag-biotin/streptavidin-PE staining and is represented in separated graphs with corresponding colours. Representative data from a single experiment using one mouse having been immunized against VIM-I.

<u>Alt text:</u> A) Four density plots and two distribution histograms characterizing size, granularity and four surface markers expression of spleen cells from an immunized mouse, revealing 3 populations. Three additional contour plots for cell surface antigen-specific or non-specific IgG expression levels of the 3 identified populations.

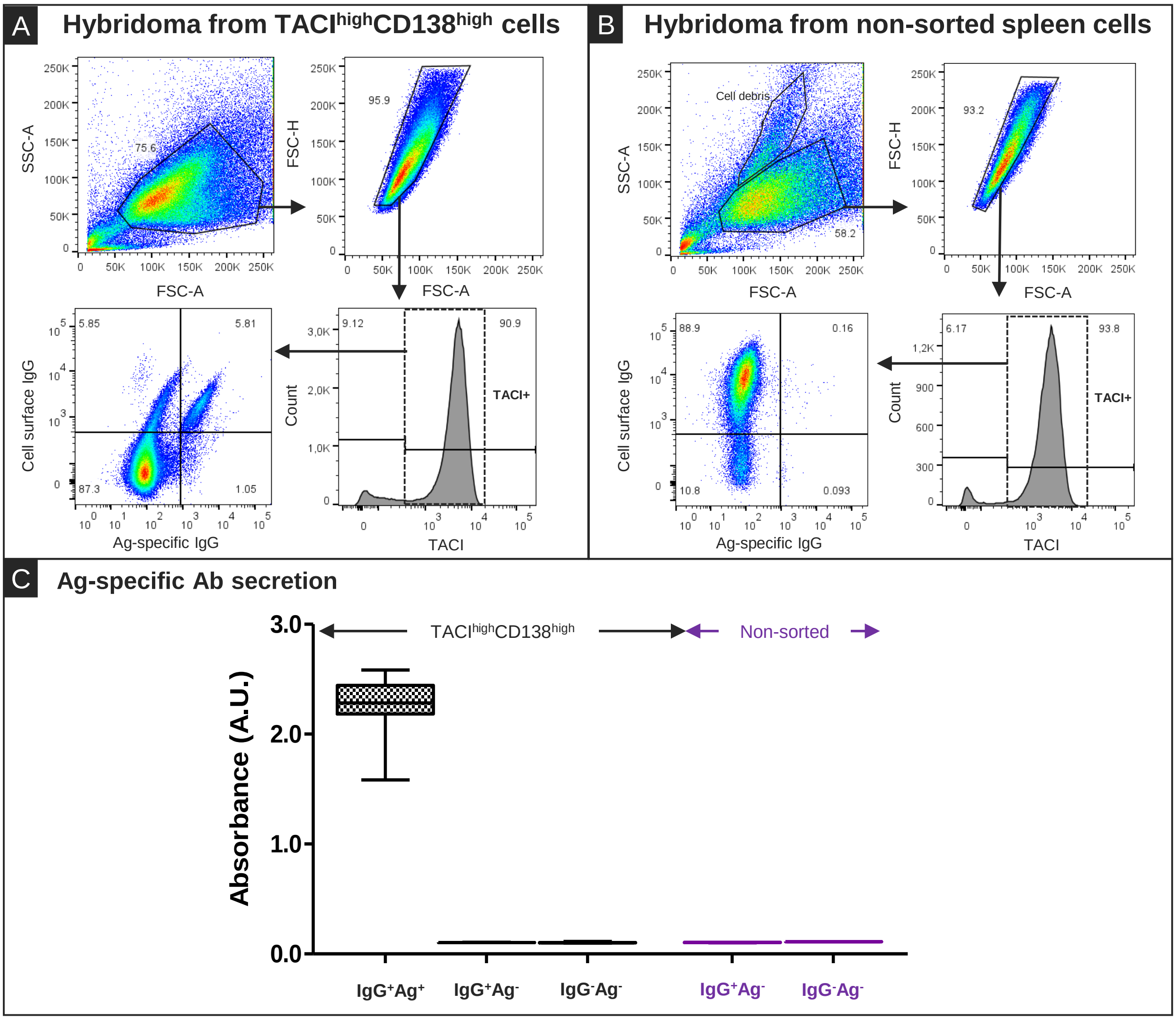


**<u>Figure S6:</u> Identification of hybridoma secreting Ag-specific antibodies after targeted fusion of TACI$^{high}$CD138$^{high}$ spleen cells.**
TACI$^{high}$CD138$^{high}$ cells from a mouse immunised against VIM were sorted by FACS and electrofused with NS1 myeloma cells. As a control. non sorted spleen cells from the same mouse were electrofused with NS1 myeloma cells. After thirteen days of culture in selective media. cells were collected and stained using TACI-APC. B220-VioBlue. and MHC-II-PerCP-Vio700. IgG-PEvio770 antibodies and Ag-biotin/Streptavidin-PE. For both TACI$^{high}$CD138$^{high}$ fused cells (A) and non sorted fused cells (B). a first gate was set on single viable hybdridoma cells through physical parameters (FSC-A vs. SSC-A) and on (FSC-H vs. FSC-A). Then. we excluded the TACII$^{-}$ cells to analyse the expression of cell surface IgG and their Ag specificity. Each of the 3 cell subsets IgG+/Ag+. IgG+/Ag-. IgG-/Ag- of TACI$^{high}$CD138$^{high}$ cells and IgG+/Ag-. IgG-/Ag- of non sorted cells was isolated in single cell mode in separate wells. After 7 days. wells containing growing cells were tested for Ag specificity with dedicated immunoassays (C). Absorbance values are represented in box and whiskers plots with minimum. first quartile. median. third quartile and maximum values.

<u>Alt text</u>: A) Three density plots and one distribution histogram characterizing size, granularity and three surface markers expression of hybridoma cells obtained after fusion of spleen antibody secreting cells, revealing three populations on the basis of antigen-specific or non-specific surface IgG expression. B) Similar plots and histograms as in panel A, but using hybridoma cells obtained after fusion with random spleen cells, revealing only two populations based on the same surface markers. C) Box plots comparing antigen-specific antibody secretion levels from the five isolated cell populations described in panels A and B. One out of five cell populations secrete antibodies.
